# Decoding the fundamental drivers of phylodynamic inference

**DOI:** 10.1101/2022.06.07.495205

**Authors:** Leo A. Featherstone, Sebastian Duchene, Timothy G. Vaughan

## Abstract

Despite its increasing role in the understanding of infectious disease transmission at the applied and theoretical levels, phylodynamics lacks a well-defined notion of ideal data and optimal sampling. We introduce a formal method to visualise and quantify the relative impact of pathogen genome sequence and sampling times—two fundamental sources of data for phylodynamics under birth-death-sampling models—to understand how each drive phylodynamic inference. Applying our method to simulations and outbreaks of SARS-CoV-2 and H1N1 Influenza data, we use this insight to elucidate fundamental trade-offs and guidelines for phylodynamic analyses to draw the most from sequence data. Phylodynamics promises to be a staple of future responses to infectious disease threats globally. Continuing research into the inherent requirements and trade-offs of phylodynamic data and inference will help ensure phylodynamic tools are wielded in ever more targeted and efficient ways.

## Introduction

Phylodynamics combines phylogenetic and epidemiological modelling to infer epidemiological dynamics from pathogen genome data (du Plessis and Stadler, 2015; Baele et al., 2018; Volz et al., 2013). Analyses are usually conducted within a Bayesian framework, meaning that the output comprises posterior distributions for parameters of interest, such as the basic reproductive number, *R*_0_ (i.e. the average number of secondary infections from the index case in an otherwise fully susceptible population). Input data usually consists of genome sequences and associated sample collection dates. In the case of birth-death-sampling models (Stadler, 2010), both sequence and date data inform the branching of inferred trees by either temporally clustering lineages or via sequence similarity. Internal nodes are assumed to co-occur with transmission events, such that they provide information about patterns of transmission that sampling time data alone cannot (Featherstone et al., 2022). Sampling times, or date data, are similar to standard epidemiological time series data while sequence data introduce the evolutionary aspect. The widely used birth-death model uses sampling times to infer a sampling rate which is also informative about transmission rates (Boskova et al., 2018; Stadler et al., 2012).

Application of phylodynamic methods has intensified since the onset of the SARS-CoV-2 pandemic. Morover, larger and more densely sequenced outbreaks are being studied. While the value of pathogen genome data is now well established, an increasingly pertinent question is whether inclusion of more sequence data after a point is of diminishing returns for some densely sequenced outbreaks (Porter et al., 2022; Hill et al., 2021). Since the answer to this question will naturally vary with each dataset and pathogen considered, we believe it is most appropriately addressed by a *method* to quantify the individual effects of date and sequence data. This would help broaden our understanding of the phylodynamic tools that now feature in infectious disease surveillance. Such a method also has the potential to help target sampling efforts of future outbreaks for optimisation of knowledge gain against resource expenditure.

Earlier work showed that sequence sampling times, referred to here as ‘date data’, can drive epidemiological inference under the birth-death model (Volz and Frost, 2014; Boskova et al., 2018; Featherstone et al., 2021). However, each stopped short of proposing a transferable way to measure this effect in regular application. The birth-death model is most applicable to the question at hand since it includes a rate of sampling. The coalescent is another a key phylodynamic model, but it typically conditions on sampling dates which therefore precluding a comparison of date and sequence effects. Some coalescent formulations include a sampling rate (Volz and Frost, 2014), however these are used less often than the birth death or standard Kingman coalescent (Kingman, 1977). The Kingman coalescent also assumes a low sampling proportion relative to population size such that its typical formulation would be inappropriate for many densely sequenced outbreaks (Boskova et al., 2018), where the question of the effect of large amounts of sequence data is most relevant.

Building upon these earlier results, we introduce a theoretical framework and a new method to quantify and visualise the effect of sequence and dates for any parameter under the birth death with continuous sampling. We focus on continuous sampling because it is most relevant to how emerging outbreak data are collected. It also classifies which data source is driving the inference, but crucially also indicates whether a binary classification is meaningful. We believe these observations will form a critical addition the phylodynamic toolkit used to inform public health decisions because they clearly quantify the added-knowledge acquired from genomes in a given analysis.

## New Methods

### Isolating date and sequence data

We conduct four analyses for a given dataset to contrast the effects of complete data, date data, sequence data, and the absence of both (fig 1A). We focus on inferring *R*_0_, with all other parameters fixed (the sampling proportion, becoming-uninfectious rate, molecular clock rate, and substitution model parameters), but this new approach is applicable to any parameter under the birth-death with any combination of priors. First, we use complete data to fit a birth-death model and infer the posterior distribution of *R*_0_. This represents the combined effects of dates and sequences. Second, to isolate the effect of date data, we remove sequence information and retain dates, thus integrating over the prior on tree topology. This is traditionally referred to as ’sampling from the prior’, but this term should be avoided in the context of models where the sampling times are treated as data, such as the birth-death. Third, to isolate the effect of sequence data, we keep sequence data and remove dates. This requires estimation of all sampling dates, analogously to how removing sequence data causes integration over topology. We use a novel Markov chain Monte Carlo (MCMC) operator to estimate dates which is implemented in the feast v17 package for BEAST 2 (Bouckaert et al., 2019). Briefly, the operator adjusts the time between the final sample date and end of the birth-death process such that dates can rescale relative to each other and in absolute time. See Fig S1 for a visual representation. Lastly, and for completeness, we conduct the analysis with both date and sequence data removed. This formally corresponds to the marginal prior conditioned on the number of samples. The resulting Wasserstein metric, *W*_*N*_, is useful for quantifying whether full data offer information in addition to the prior.

**Figure 1:**
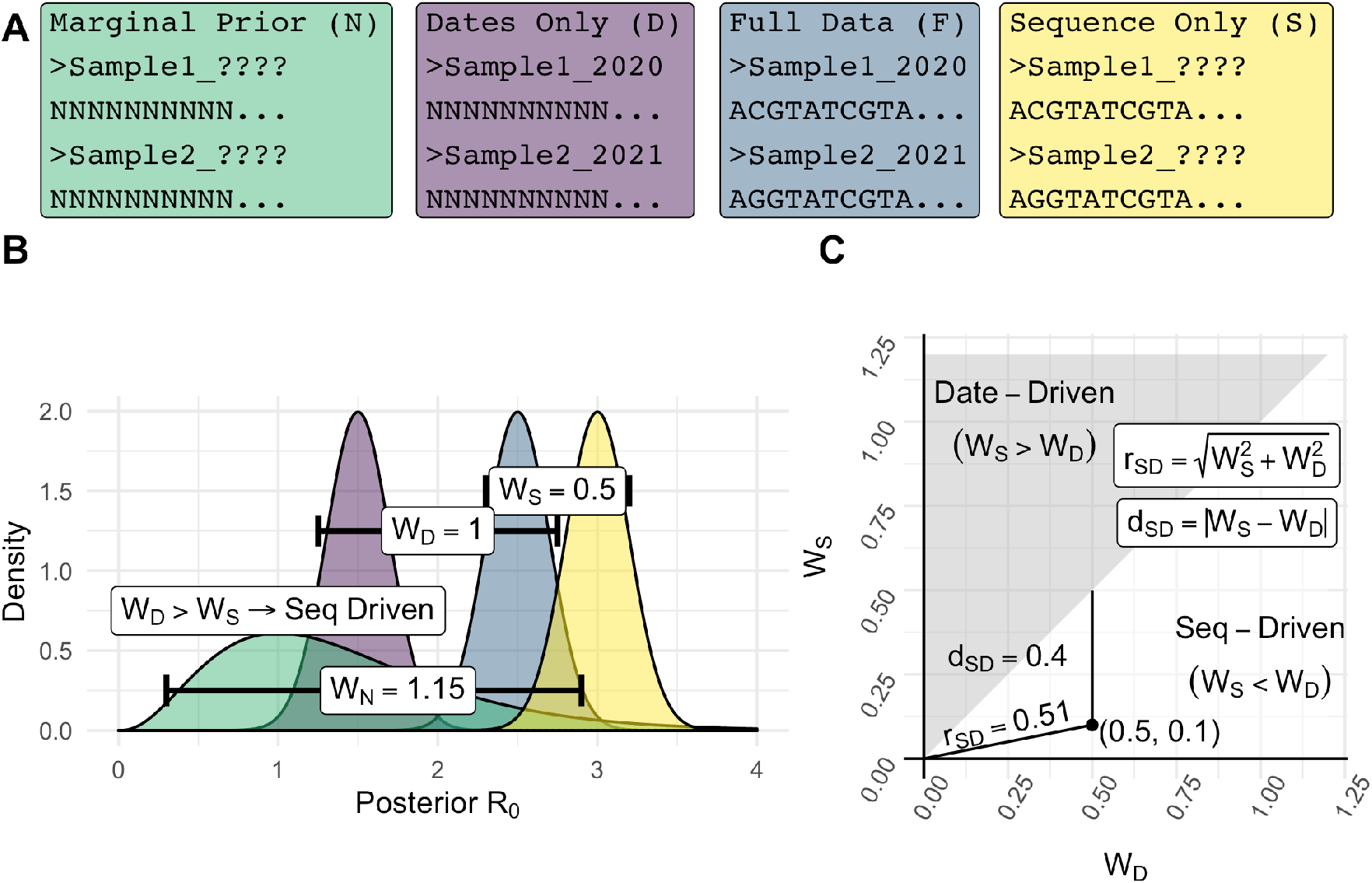
Graphical summary of the process to quantify signal and classify signal drivers. **A**) Coloured boxes give examples of data under each of the 4 treatments with letters in brackets giving shorthand notation for each. From left to right: *Marginal Prior* results from the removal of both date and sequence data. *Dates Only* includes date data while ignoring sequence data through a constant phylogenetic likelihood. This can be represented as converting all sequence characters to ‘N’ (i.e. alignments of entirely missing data). *Full data* represents the usual combination of both sequence and date data. This produces a reference distribution from which the Wasserstein metric to other posteriors is calculated. *Sequence Only* corresponds to the removal and re-estimation of dates while sequence data is retained. **B**) Example posterior output for *R*_0_ with colour corresponding to each treatment in **A**. The Wasserstein metric is calculated as difference in inverse distribution function of each posterior from *Full Data* integrated over 0 to 1. Example values for the Wasserstein metric are given in white boxes. **C**) The plane with x and y axes *W*_*D*_ and *W*_*S*_ and shaded classification regions. *r*_*SD*_ is the Euclidean distance from the origin to a point (*W*_*D*_, *W*_*S*_), with higher values indicating that one or both of data and sequence data drive differing signals from the reference posterior. *d*_*SD*_ is the vertical distance from a point (*W*_*D*_, *W*_*S*_) to the line *y* = *x*, with points closer to this line corresponding to more similar data and sequence data effects such that classification is less meaningful. In the example, distance from the posterior under only sequence data to the full data posterior (*W*_*S*_) is smallest, leading to classification as ‘Seq-Driven’.

### Quantifying data signal

We employ the Wasserstein metric in one dimension to measure a “distance” between each of the sequence posterior, date posterior, or marginal prior, and the posterior derived from the complete data. We write these distances as *W*_*•*_, with *•* being *D, S*, or *N* for the sequence, date, and marginal prior distributions, respectively. For example, the Wasserstein distance *W*_*D*_ from the date-data to complete data posterior is:

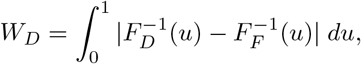

where *F*_*D*_ and *F*_*F*_ are cumulative distribution functions for the posterior *R*_0_ under date and complete data respectively. The units of *W*_*•*_ are equivalent to the units of the parameter of interest (e.g. years^*−*1^ for the birth rate, becoming un-infectious rate, sampling rate, and unitless for *R*_0_). See Fig S2 for a visual interpretation of the expression. We used the transport R package to calculate the Wasserstein metric.

As in Fig 1C, we can now consider a plane where the axes are *W*_*D*_ and *W*_*S*_. We classify the data source with the lowest Wasserstein distance from the complete data posterior as contributing most to the posterior from full data. We refer to this as a classifier, although we emphasise that this not a classifier as in machine learning literature. There is no prediction or statistical modelling of the driving data source beyond *min*(*W*_*D*_, *W*_*S*_). In this case the lines *y* = *x* marks the classification boundary as in the shading in Fig 1C.

Finally, we can quantify the disagreement in signal between each data source. We define disagreement with respect to the full data posterior, *r*_*SD*_ as the magnitude of the vector 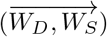 leading to each point in the plane. This is also the radius from the origin to the point. Values near zero indicate that the posteriors under date data, sequence, and complete data are all near identical and classification as date- or sequence-driven is less meaningful. Larger values signify that one or both data sources drive differing posteriors and classification is more meaningful. We also define disagreement without respect to full data *d*_*SD*_ = |*W*_*S*_ *− W*_*D*_| as a quantification of disagreement between date and sequence posteriors without reference to the full data (Fig S3C). Visually, this corresponds to the vertical distance to the classification boundary (*y* = *x*) such that smaller values correspond to less meaningful classification. *r*_*SD*_ and *d*_*SD*_ are similar in that when *r*_*SD*_ is near-zero, *d*_*SD*_ necessarily is too. *d*_*SD*_ also accounts for the case where *r*_*SD*_ is high, but both date and sequence data have similarly sized effects. In this case, *r*_*SD*_ is higher while *d*_*SD*_ is lower and classification of one or another as driving analysis is inaccurate.

## Results

We simulated 600 alignments to explore the differing signals in date and sequence data using the Wasserstein metric. These derive from 100 simulated outbreaks of 500 cases, sampled with proportion 1, 0.5, or 0.05 (*n* = 25, 250, or 500 accordingly) and used to simulate sequences with an evolutionary rate of 10^*−*3^ or 10^*−*5^ (subs/site/time). Higher evolutionary rates induce more site patterns and therefore more informative sequence data. We estimated *R*_0_ under each data treatment with all other parameters fixed using a birth-death tree prior. In all analyses, simulated data provided information in addition to the prior (*W*_*N*_ *>* 0.06, Fig S5). Among the 600 datasets, we observe a mixture of cases where date and sequence data infer similar or dissimilar posterior *R*_0_. This supports the core assumption that date and sequence data can have differing signals concealed in their combination (Fig 2). Classification using the Wasserstein metric results in a mix of date and sequence driven classifications, supporting that our proposed method is sensitive to differences between datasets (Fig 3). Most datasets were classified as date-driven (372/600), which is consistent with earlier work showing that dates are highly influential under the birth-death (Volz and Frost, 2014).

**Figure 2:**
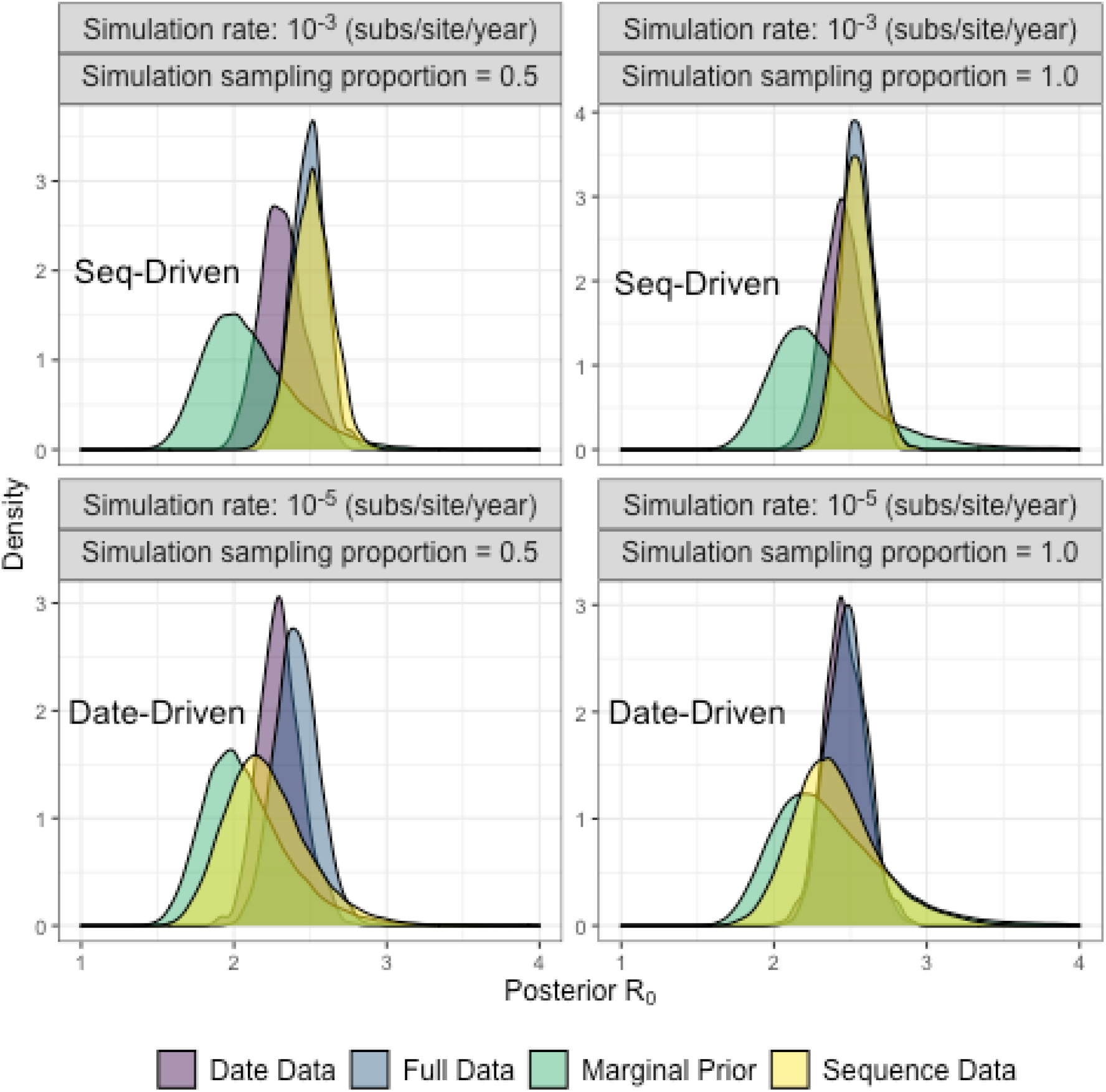
Comparison of full data, date data, sequence data, and no data posterior *R*_0_ for four of the 600 simulated datasets.

**Figure 3:**
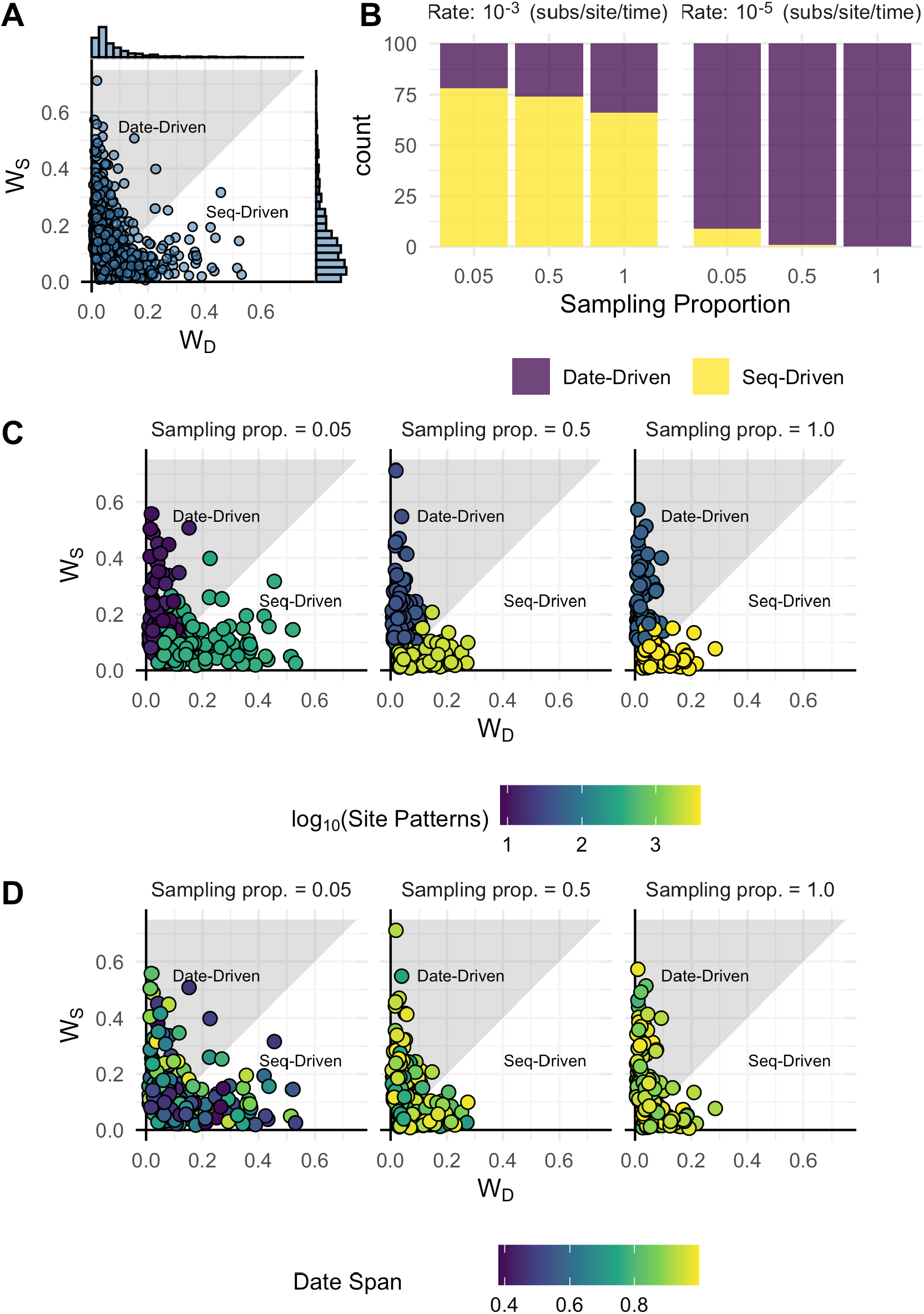
**A**) Each point represents (*W*_*D*_, *W*_*S*_) for one of the 600 simulated datasets with marginal histograms corresponding to the distribution of *W*_*D*_ and *W*_*S*_ respectively. **B**) The number of simulated datasets classified as Date of Seq Driven, stratified by evolutionary rate and sampling proportion. **C**) Points coloured by number of site patterns. Lower site patterns tends to co-occur with date-driven classification. **D**) Points coloured by date span with no clear patterns corresponding with classification as for site patterns.

### Reliability of Wasserstein Metric

We conducted a subsampling analysis to ensure that Wasserstein metric values reflected differences between date and sequence data, rather than noise alone. For each of the 600 simulated datasets, we subsampled posterior *R*_0_ distributions corresponding to each of the 4 data treatments 100 times, taking a number of samples equal to half the length of the post-burnin posterior. This yielded 60,000 subsampled posteriors for which we re-calcualted Wasserstein values and reclassified each as in the original simulation study. Of the 60,000, subsampled posteriors, only 99 were misclassified across 99 of the 600 original datasets. In other words, misclassification occurred only once out of 100 replicates for 99 of the 600 simulated datasets.

Of the 600 simulated datasets, those where misclassification occurred had substantially smaller differences between Wasserstein distance to the date-only and sequence-only posteriors (*d*_*SD*_) (Figure S3). Misclassification occurred where *d*_*SD*_ *≤* 8.27*×*10^*−*2^ in the complete *R*_0_ posteriors, with no error above this level. Differences below 8.27 *×* 10^*−*2^ correspond to a level of difference between data and sequence only posteriors where classification is of little significance (i.e. *R*_0_ *±* 0.08 assuming both posteriors have similar uncertainty). Classification is wholly reliable above this threshold, validating the trends observed in the simulation study and that the Wasserstein metric is sensitive to differing signals concealed in date and sequence data.

### Observations about the effects of Sequence and Dates

The distribution of *W*_*S*_ is more diffuse than *W*_*D*_, meaning the sequence data posterior tends to differ more from full data than date data (Fig 3 A). This again aligns with previous results showing that date data usually drive inference under the birth-death.

Low sequence diversity, measured here in the number of site patterns, seems to preclude sequence data from driving inference (Fig 3C, Fig S4). This matches the expectation that fewer site patterns results in less sequence information to inform the posterior. It appears as a necessary, but not sufficient condition for a dataset to have at least one site pattern per sample in order for analysis to be sequence driven (Fig S6). On the other hand, the date span does not follow an equivalent trend with lower diversity coinciding with analyses being sequence driven. Here the relative date span is the time between the first and last sample, divided by the height of the outbreak tree and thus should be indicative of the information content of the dates through the proportion the outbreak’s duration that they capture. The distribution of relative date-span appears random across classifications, unlike the distribution of site patterns (Fig 3C, Fig S4).

Sampling proportion appears to play a secondary role in driving the influence of sequence data (Fig 3B, animated [here](https://github.com/LeoFeatherstone/phyloDataSignal/blob/main/figures/wassTraj.gif)). Lower sampling proportions coincide with a higher relative proportion of sequence-driven datasets. This matches expectation because lower sampling proportion increases the likelihood of sampling more divergent lineages, resulting in more site patterns to drive inference.

One of the core assumptions of phylodynamics is that inferred phylogenetic trees bear a resemblance in their shape and timing to underlying epidemiological networks (Featherstone et al., 2022). Based on this, we would expect that analyses with less uncertainty in posterior epidemiological estimates have posterior tree distributions that are more divergent. To test this expectation, we calculated tree distance metrics pairwise for each of the 2400 posterior tree distributions, originating from the 600 simulated outbreaks and 4 data treatments. We sampled 100 trees from each posterior and considered the pairwise topological distance between each. We focus on the mutual clulstering information (MCI) introduced by Smith (2020) (Fig S7), but all observed patterns were repeated when using other metrics including Robinson-Foulds and Information-corrected Robinson-Foulds. The full data treatment consistently had the lowest pairwise MCI since these analyses incorporated the most information to constrain tree-space. The marginal prior, reflecting the least information, reported the highest MCI values. Between these two extremes, sequence-only analyses consistently had a lower MCI than date-only treatments. This can be explained by sequence data providing information about favoured topologies via the phylogenetic likelihood. Conversely, dateonly data incorporate no topological information and consistently have higher MCI values. Given that date-data tend to drive inference, especially where evolutionary rate is low and to an extent where sampling proportion is high, this presents strong evidence that the *chronology* of tips can be equally, if not more informative than the *topology* of trees. This highlights the importance of curating sampling time data as carefully as sequence data is curated. Moreover, this applies at the phylogenetic level of phylodynamic inference, as distinct from the choice of phylodynamic model and its parameters.

### Empirical Results

#### Australian SARS-CoV-2 Clusters

We analysed data from two SARS-CoV-2 transmission clusters from 2020 in Australia to demonstrate that date-driven and sequence-driven analyses can arise in practice (table 1). These clusters were chosen out of the 595 identified by Lane et al. (2021) because they possessed the most complete sampling-time data to compare with sequence data. They are also ideal examples of the size and type of genome dataset that public health practitioners may seek phylodynamic insight from as each sample most likely reflects transmission stemming from a single, unknown source as supported by contact tracing.

**Table 1:** SARS-CoV-2 Clusters

|  | Cluster 1 | Cluster 2 |
| --- | --- | --- |
| n | 112 | 188 |
| Classification | Seq-Driven | Date-Driven |
| $r_{SD}$ | 0.223 | 0.009 |
| $d_{SD}$ | 0.078 | 0.008 |
| $W_D$ | 0.192 | 0.001 |
| $W_S$ | 0.114 | 0.009 |
| $W_N$ | 0.325 | 0.481 |

Analysis of the first cluster is classified as sequence-driven, with *r*_*SD*_ = 0.223 indicating an appreciable difference between the sequence posterior and complete data. *d*_*SD*_ = 0.078 adds that date data also drives an effect of similar size, offering the interpretation that both date and sequence data are influential in this analysis. The second cluster is classified as date-driven, but with *r*_*SD*_ = 0.009 and *d*_*SD*_ = 0.008. The low *r*_*SD*_ value indicates a near-negligible difference between date, sequence, and complete data posteriors. Since *r*_*SD*_ is low, *d*_*SD*_ is necessarily also low. Due to this, classification is effectively meaningless and it can be concluded that both date and sequence data drive a highly congruent signal. Moreover, *W*_*N*_ for both analyses is more than double each of *W*_*S*_ and *W*_*D*_, which affirms that both sources of data contribute to the posterior deviating from the prior and are therefore informative with respect to the prior in both analyses.

#### 2009 H1N1

The above simulation study and SARS-CoV-2 Empirical data consist of analyses where all parameters are fixed except *R*_0_. However, this extent of prior certainty is rare in practice. In particular, there exists an inherent trade-off between prior certainty in the evolutionary rate and the strength of signal due to sequence data. Moreover, this affects inference of the sampling proportion, which relies on sequence information to inform the evolutionary distance between samples and tMRCA, in turn informing the proportion of lineages captured in sampling. To explore the effects of these trade-offs, we consider North-American H1N1 data from the 2009 Swine-Flu pandemic originally studied by Hedge et al. (2013). This dataset was chose because it represents a small sample from a much larger outbreak, which helps to ensure to ensure the presence of sequence information with which to test the effects of prior uncertainty in the evolutionary rate.

We again fit a birth-death process, estimating two *R*_*e*_ values before and after early July 2009 (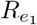 and 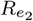 respectively) and the sampling proportion (*p*). Analyses were conducted with either a fixed evolutionary rate, 4 *×* 10^*−*^3 subs/site/year following Hedge et al. (2013), or a uniform prior U(10^*−*4^, 10^*−*2^), and under each of the four data treatments defined above. We placed a non-informative *β*(1, 1) prior on the sampling proportion and fixed the becoming-uninfectious rate 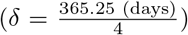, such that comparing analyses with a fixed or uniform prior on the evolutionary rate reveals the effects of uncertainty in evolutionary rate on *R*_*e*_ and sampling proportion. Posterior distributions for *R*_*e*_ and *p* are presented in Fig 4 and Wasserstein values in table 2.

**Table 2:**
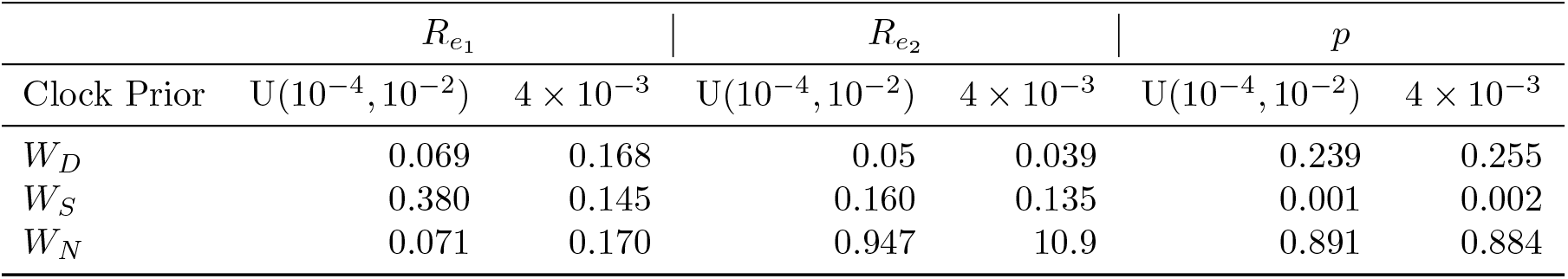
2009 H1N1

**Table 3:** Practical considerations for inference stratified by relevance to phylogenetic inference or the phylodynamic model (Birth-Death here).

| Trade-off / Un-identifiability / Challenge in inference | Mitigating Action | Effect |
| --- | --- | --- |
| **Phylogenetic Level** |  |  |
| Ideally $\geq 1$ Site Patterns per Sample | Sample population less frequently, if possible, for longer. | More site patterns |
| **Phylodynamic model level (Birth-Death)** |  |  |
| tMRCA $\leftrightarrow r^\dagger$ | Reducing prior uncertainty in one reduces posterior uncertainty in the other | Reduces mutual un-identifiability |
| Un-identifiability of $R_0, \delta, p$ | Fix at least one parameter (Louca et al., 2021) | Localise inference to correct congruence class of parameters |
| $R_0 \leftrightarrow r$ | Narrowing prior $r$ | More sequence information available to ascertain $R_0$ |
| $p^\ddagger \leftrightarrow r$ | | More sequence information available to ascertain $p$ |
| Site Patterns $\rightarrow p^\ddagger$ | Consider if fewer site patterns in sample is inflating posterior $p$ erroneously | |

<sup>†</sup> The clock rate in subs/site/year. <sup>‡</sup> Effect of sequence data much stronger than uncertainty in prior $r$ .

**Figure 4:**
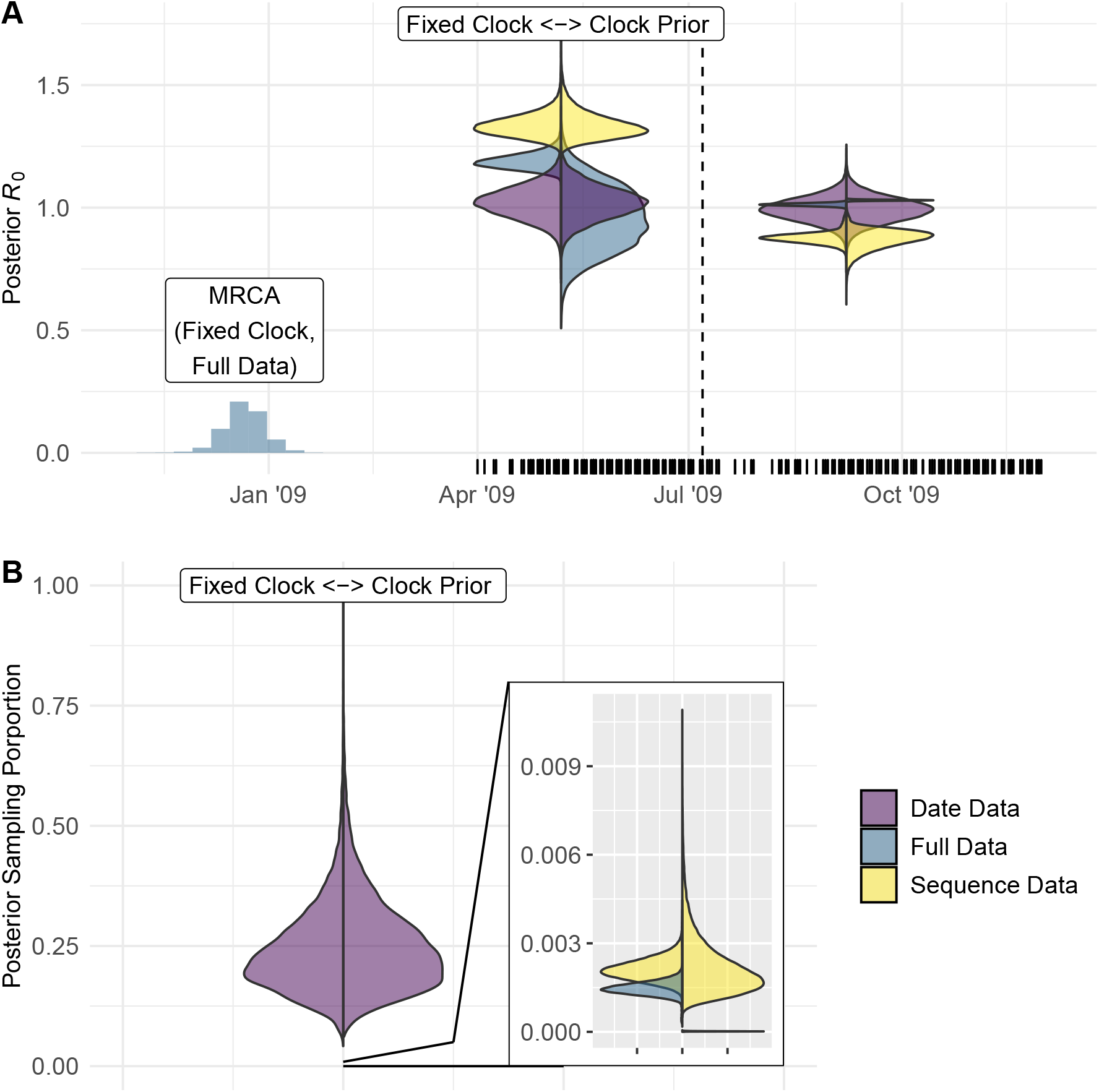
Posterior estimates for *R*_0_ and *p* for the 2009 H1N1 dataset. **A)** Histogram gives shape of posterior density for the tMRCA from analysis of the full data under a strict clock. Counts are re-scaled by a factor of 10^*−*5^. Vertical marks along the date axis represent sampling times in the dataset. The vertical dashed line marks the change time between 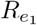 and 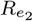. Asymmetrical violin plots give the posterior density for *R*_*e*_ under each data treatment with the marginal prior omitted. Distributions on the left correspond to analyses with a fixed evolutionary rate and those on the right correspond analyses with a uniform prior on the evolutionary rate. The more diffuse clock prior decreases the influence of date data. **B)** An asymmetrical violin plot of the posterior sampling proportion with the left and right corresponding to fixed and uniform clock priors identically to **A**. Sequence data drives inference for both clock prior configurations.

The patterns in posterior *R*_*e*_ under each treatment verify the trade-off between estimating tip dates and evolutionary rate. This is reflected in higher *W*_*S*_ values when using a prior on the evolutionary rate than when fixing it (Table 2), which corresponds to posterior *R*_*e*_ under Full Data overlapping less with the posterior from sequence data. Moreover, the sequence data posterior is wider where the uniform clock prior is used, reflecting that more sequence information is devoted to ascertaining the evolutionary rate where the prior is more diffuse, which would otherwise further ascertain *R*_*e*_ (Fig. 4 A). There is also a disparity between the posterior tMRCA between each clock prior under Full Data (Only the fixed prior tMRCA is shown in Fig 4 A), which is expected given the well known non-identifiability between the evolutionary rate and tMRCA.

Estimates of sampling proportion appear to be strongly sequence driven regardless of evolutionary rate prior (Fig 4 B). The posterior sampling proportion is two orders of magnitude lower for the sequence data and full date treatments, both containing sequence information, than for date data alone. These estimates are far more in line with expectation too, given the disparity between the H1N1 sample size (n = 100), and scale of the 2009 H1N1 pandemic. This is expected because the diffuse *β*(1, 1) prior placed on *p* made sequence information essential to informing the evolutionary distance between samples and driving estimates of sampling proportion lower. Conversely, estimates with date-only data are much higher and more uncertain owing to the lack of sequence information. The evolutionary rate prior drives a similar trend as for *R*_0_, but its effect is diminished by the major effect of sequence data. Altogether, this highlights that different parameters can be draw from date and sequence data to different extents.

We do not present the marginal prior data in Fig 4 because each corresponding posterior is un-informatively broad (i.e. effectively uniform over the prior domain). However, this too supports our assertion of a trade-off between sequence information and posterior certainty in sampling proportion because of its contrast to the simulation studies and SARS-CoV-2 datasets where sampling proportion was fixed. The only information included in the marginal prior treatment is the sample size, and this is most informative when the sampling proportion is fixed. However, in the H1N1 analyses, the diffuse prior on sampling proportion means sample size alone is insufficient to drive an informatively narrow posterior and sequence is the primary driver of inference.

Taken together, the results of the H1N1 dataset demonstrate that uncertainty in evolutionary rate prior partly determines the strength of effect for sequence data, since any information in sequence data is used in shifting from prior to posterior. In addition, sampling proportion demands sufficient sequence information for inference, and this cannot be substituted by sampling times as readily as has been shown for *R*_*e*_ Featherstone et al. (2021); Volz and Frost (2014).

## Discussion

The results of our simulation study add clarity practical implications to previous results showing that sampling times contribute substantially to phylodynamic inference under the birth death (Volz and Frost, 2014; Featherstone et al., 2021). We also demonstrate that sequence data are not always secondary in influence and can drive inference of *R*_0_ in some instances, affirming the sensitivity of the birth-death to the signal encoded in sequence data. We demonstrate that the number of site patterns is key enabling sequence-driven inference, but this is not the sole factor. Over-sampling from a pathogen population can also dilute sequence information and pivot analyses to being date driven.

The tendency for date data to drive inference as site patterns decrease and sampling proportion increases can be explained by the reduction in uncertainty that each data source offers. Dates impose hard bounds by restricting tree space to a subset of trees that agree with the chronology of sampling times. Conversely, sequence data inform topology through phylogenetic likelihood, but do not definitively constrain tree space in the same way as date data. The net result is that sequence data must explore a larger tree-space than date data. Thus, as the information in sequence decreases, such as though intensified sampling lowering the number of site patterns per sample, the disparity between the sequence and date data tree spaces increases and dates more often drive analyses. See Fig S8 for visual explanation.

We note that having sufficient sequence information to drive inference is a related but ultimately different concept to the phylodynamic threshold, which is defined as the sampling timespan needed for a pathogen to display temporal signal (Duchene et al., 2020). Our analyses show that surpassing the phyloydnamic threshold (i.e. having temporal signal present in the data) is generally not a sufficient condition for sequence-driven inference. Conversely though, we demonstrate with the H1N1 dataset that increased prior certainty in the evolutionary rate increases the effect of sequence data in analysis. This means that an assessment of the phylodynamic threshold is highly useful in practice to promote sequence-driven inference, but only retrospective analysis using the methods introduced here can provide conclusive proof about the contribution of the sequence data under consideration.

### Practical Implications

The central message of our results is that sampling time data increasingly drive phylodynamic analyses as datasets grow in the proportion of sampled cases. While information from sample times can be a boon to phylodynamic analyses when the sampling process is properly modeled, dependence on this information can become a vulnerability as sampling processes deviate from the assumptions of phylodynamic models over time and space. Thus, the phylodynamic-value of a dataset must take into consideration the amount of sequence and date information available to direct inference over tree space, rather than the size of the dataset alone. — Bigger is not automatically better. To this end we recommend a cautious approach to sampling, with the recognition that each additional sample is increasingly costly to the analysis in requiring sequence data to navigate a disproportionately large tree space in comparison to sampling times. Therefore, samples that are both identical in sampling time and sequence, such as from a super spreading event, should be avoided. As a basic start point, having at least one site pattern per sample greatly increases the likelihood of a sequence driven analysis (Fig. S6).

Huge databases of pathogen sequence data, such as GISAID and Nextstrain are immense assets to the phylodyanmcis research program, but careful curation of datasets with respect to both sampling times and sequence quality are essential to providing valuable phylodynamic results. Just as it is valuable to surpass the phylodynamic threshold in sampling, it is of equal value *not* to surpass the notional threshold of redundancy in sampling from phylodynamic studies.

Moreover, different parameters require date and sequence information for accurate inference to differing extents. The ability to infer sampling proportion is integral to the value of phylodynamics as it allows for estimation of how many unknown cases there may be in an outbreak. We show sampling proportion is highly sequence driven, which necessitates both sampling to maximise sequence information relative to date information, and measurement of the effect of sequence data in analysis to comment on the reliability of estimated sampling proportions.

Fixing or constraining parameters in analyses where it is justifiable is also critical to ensuring sequence information surfaces in posterior results. For example, fixing the evolutionary rate is ideal if possible. Providing topological constraints between samples would also help concentrate sequence information in the data by reducing the tree space analyses must traverse. These guidelines are complementary to recent work due to Louca et al. (2021), showing that at least one of the unidentifiable parameters under the birth death (*R*_*e*_, *δ*, and *p*) needs to be fixed for accurate inference from the state space of analyses (i.e. the tree and generative generative parameters).

A final practical message is that a phylodynamically ideal microbe would, for example, have an evolutionary rate of of 10^*−*1^ (subs/site/time) on a genome 10^10^ bases with consistent clocklike evolution to uniquely barcode every transmission event. Clearly, no such microbe exists and phylodyamics broadly operates over microbes with genomes between 10^4^ and 10^8^ bases in size and evolution rates from 10^*−*2^ to 10^*−*8^ (subs/site/year) (Biek et al., 2015). In this view, there will never be a perfect dataset or pathogen to analyse. The challenge instead becomes sampling and constraining analyses parsimoniously enough to ensure the epidemiological signature encoded in sequence data is returned as well as measuring this effect, such as by the methods developed here.

### Future Directions

The work presents the beginning of deeper interrogations of the drivers of phylodynamic inference. Such an understanding is critical as phylodynamics becomes a mainstay of infectious disease surveillance.

Here we have presented a method, based on the Wasserstein metric, for post-hoc analysis of the drivers of phylodynamic inference. A critical future direction would be to develop predictive methods with explicit mathematical formulae for the information content of phylodynamic data. Such a toolkit would enable a theory of optimal sampling for phylodynamic inference to be explicitly described. However, we stress that a method to assay the drivers of inference, such as the work presented here, needed to be developed first to help measure the effectiveness of any future predictive methods.

### Data Archival

All scripts used to simulate and analyse data are available at https://github.com/LeoFeatherstone/phyloDataSignal. git. The Feast package, containing our date estimator, is available at https://github.com/tgvaughan/feast.git.

## Methods

### Simulation Study

We simulated 100 outbreaks of 500 cases under a birth death process using the `Tree-Sim` R package (Stadler, 2019). The birth rate was set to 2.5, death rate 1, corresponding to *R*_0_ = 2.5. Sampling probability was set to *p* = 1, resulting in trees with 500 tips. We then extended this to a set of 300 outbreaks by sampling again with probability *p* = 0.05 and *p* = 0.5, resulting in trees of 25 and 50 tips. We used a consistent seed such that each outbreak with *p* = 0.05 or *p* = 0.5 corresponds to a subsample of another with *p* = 1, allowing us to asses the effect of sampling proportion on inferring *W*_*•*_. For each outbreak, we used Seq-Gen to sequence alignments of length 20000, which is roughly average for RNA viruses (Sanjuán et al., 2010; Rambaut and Grass, 1997). We set an Jukes-Cantor model with evolutionary rate set to either 10^*−*3^ or 10^*−*^5 subs/site/time using Seq-Gen Rambaut and Grass (1997). Our choice of evolutionary rates allows us to compare the effects higher and lower sequence information with the former corresponding to about 20 substitutions per infection, and 0.2 for the latter. The above resulted in 600 alignments to test in the four treatments described above. We analysed each under a birth-death model using BEAST v2.6 Bouckaert et al. (2019) with a Uniform[0, 5] prior for *R*_0_ and all other parameters set to the true value for simplicity and to disentangle any impacts of parameter nonidentifiability (Louca et al., 2021).

### Empirical Data

All empirical analyses were conducted using BEAST v2.6.6 (Bouckaert et al., 2019) and MCMC chains were run until all parameters of interest had ESS above 200 following burnin. The only exception is the marginal prior for the H1N1 data with a fixed clock. Achieving ESS *>* 200 for 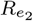 required a prohibitively long chain.

### SARS-CoV-2

We analysed two similar SARS-CoV-2 datasets taken from Lane et al. (2021). They consisted of 112 and 188 samples respectively.

We analysed each dataset under the four conditions above. In each, we placed a Lognormal(mean = 1, sd = 1.25) prior on *R*_0_ and an Inv-Gamma(*α* = 5.807, *β* = 346.020) prior on the becoming-uninfectious rate (*δ*) following estimates of the duration of infection 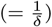 in Lauer et al. (2020). We also fixed the sampling proportion to *p* = 0.8 since every known Victorian SARS-CoV-2 case was sequenced at this stage of the pandemic, with a roughly 20% sequencing failure rate. We also placed an Exp(mean = 0.019) prior on the origin, corresponding to a lag of up to one week between the index case and the first putative transmission event.

### H1N1

We analysed North American H1N1 Swine flu samples taken from the 2009 swine flu pandemic studied by Hedge et al. (2013). We fit a birth-death model with two intervals for *R*_*e*_, before and after 2020-07-07. This date sits in a breakpoint in sampling intensity and was chosen to separate transmission dynamics between the early and late stages of the sampling timespan.

We fixed the becoming infectious rate to 91, corresponding to a duration of infection of 4 days. We also placed a *β*(1, 1) prior on the sampling proportion and used an HKY with 4 gamma categories. We compared the effect of two prior configurations for the evolutionary rate, using either a fixed rate a 4 *×* 10^*−*3^ subs/site/year or a uniform prior (U(10^*−*4^, 10^*−*2^)). All other priors were left as defaults.

---

**Figure S1:**
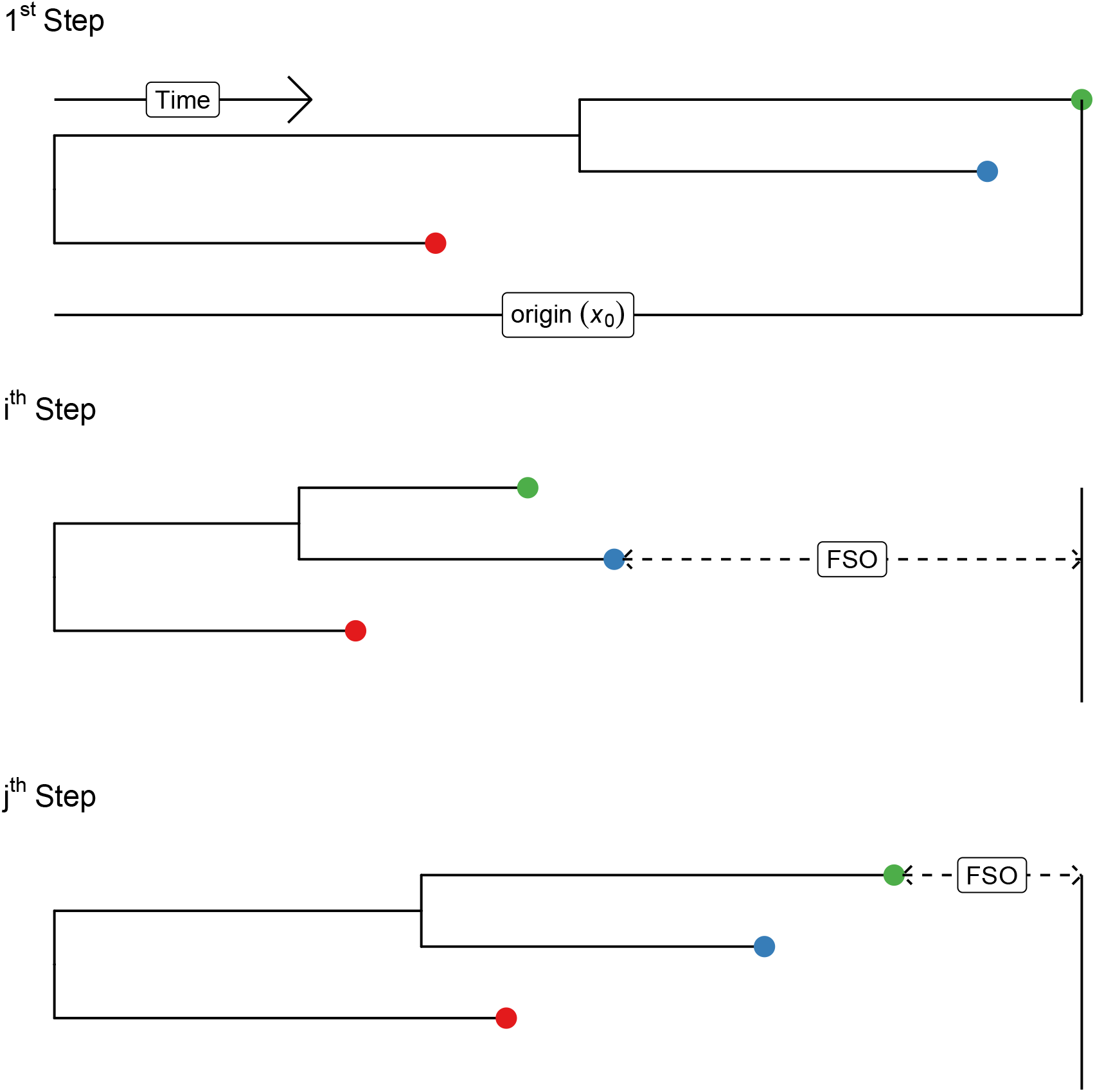
Visual intuition for the novel tip-date operator that allows for inference from sequence only datasets. Each row represents a state in an hypothetical MCMC chain. The Final Sample Offset (FSO) is the time between the final sample and the theoretical end of the proposed birth-death process. It is normally held constant in inference. The tip date operator allows for new proposals of the FSO which allows tip dates to rescale relative one each other and in absolute time. The origin time does not contain the FSO.

**Figure S2:**
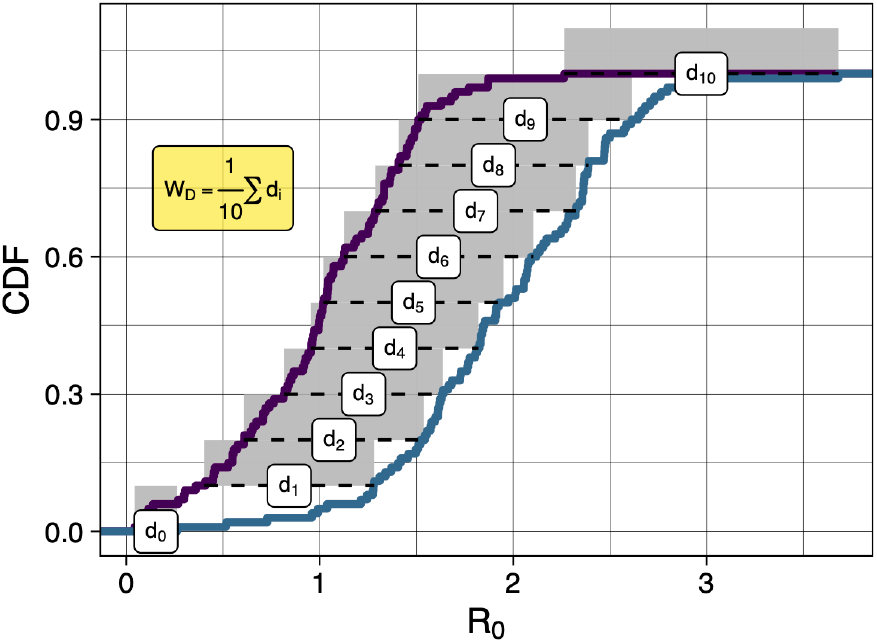
Visual intuition for calculation of the Wasserstein metric inspired by Kolouri et al. (2019). It can be thought of as integration over the horizontal distance between points of the cumulative distribution function. For two sets of posterior samples from an MCMC, the 10 horizontal distances would be replaced by the the size of the greatest set of samples.

**Figure S3:**
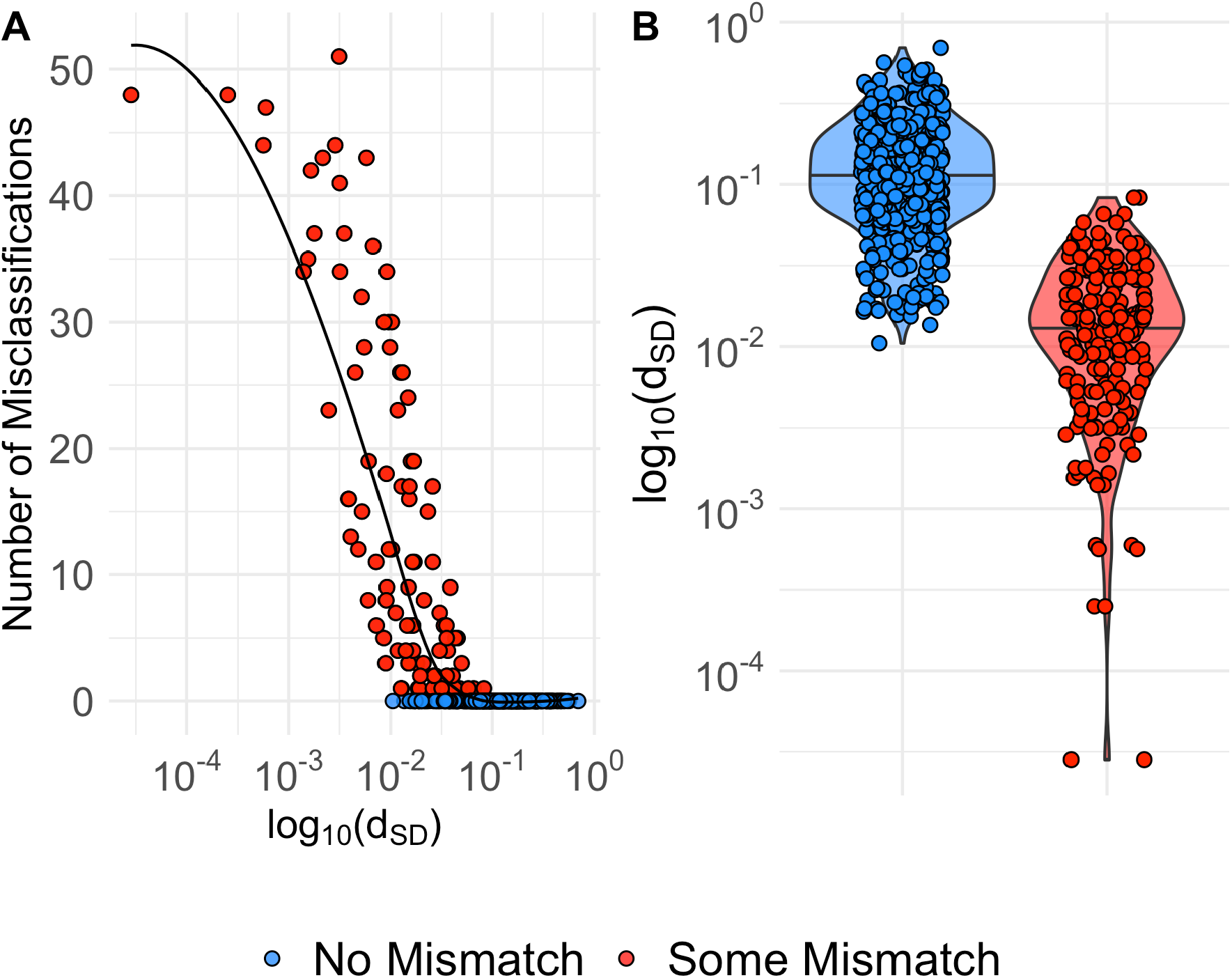
**A**) Each point presents the number of misclassification in subsampling posterior *R*_0_ for each of the 600 simulated datasets. X-axis is the log transformed difference between *W*_*S*_ and *W*_*D*_. **B**) Violin plots with jittered points of the difference between *W*_*S*_ and *W*_*D*_ for simulated alignments where there was wither some one no misclassification. Both **A** and **B** support that misclassication only occurs where the difference between *W*_*S*_ and *W*_*D*_ is negligible, and classification is not meaningful in the first instance.

**Figure S4:**
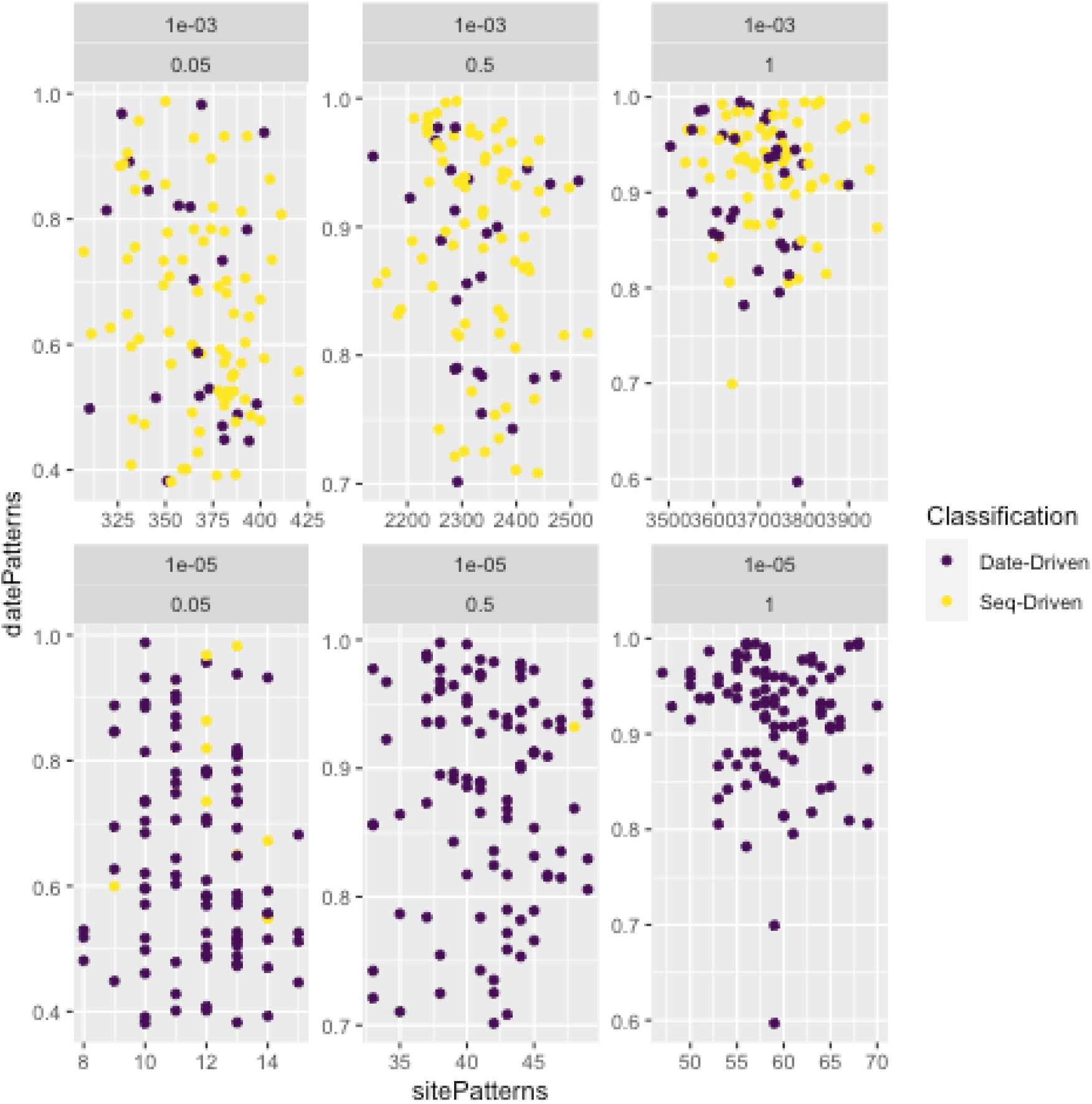
Date patterns 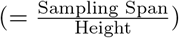 against site patterns for each simulated dataset. Plots are separated by evolutionary rate and sampling proportion such that there are 600 points coloured by Wasserstein Classification across the entire figure. Higher evolutionary rates increase the proportion of Sequence driven datasets, but within each rate there is no clear pattern in date patterns, site patterns, or sampling rate driving classification.

**Figure S5:**
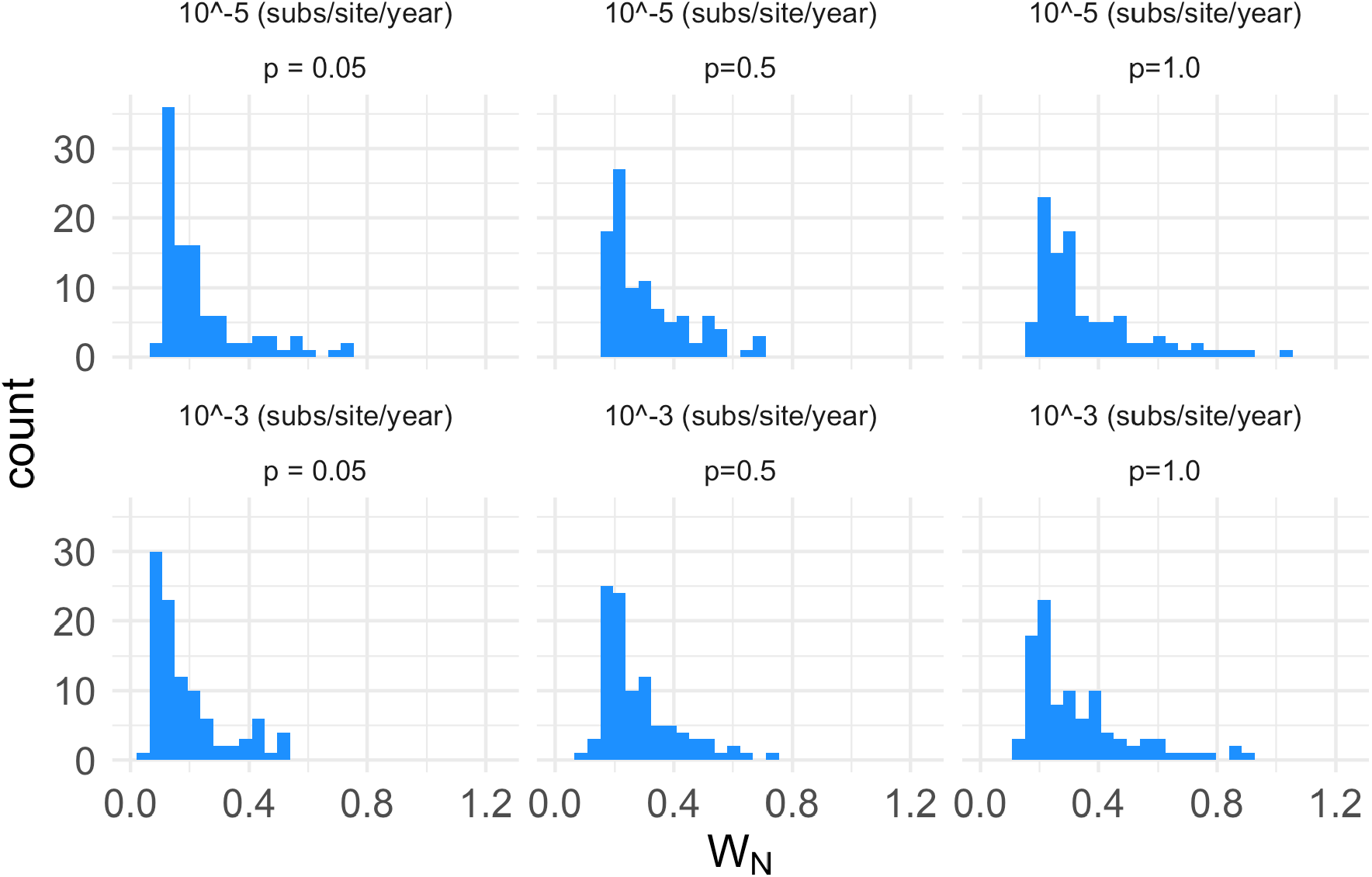
Histogram of *W*_*N*_ for each simulated dataset, separated by evolutionary rate and sampling proportion. *W*_*N*_ ranges from 0.06 to 1.02, such that simulated data provide additional information beyond that of the prior in each analysis.

**Figure S6:**
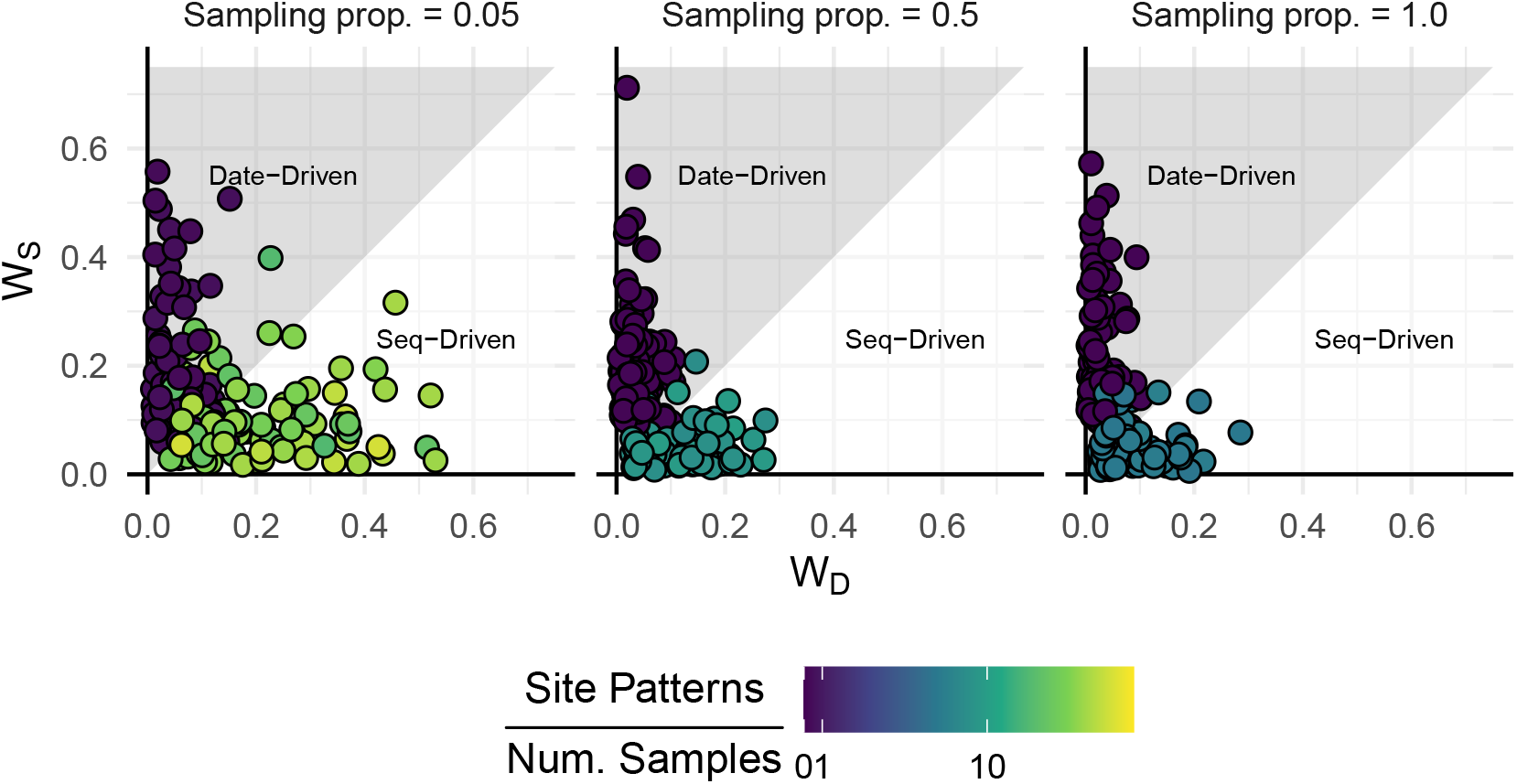
Number of site patterns per sample across each sampling proportion used in the simulation study. Analyses classified as sequence driven always have more than one site pattern per sample. This is not a sufficient condition to an analysis being sequence driven, as seen in some analyses with higher site patterns per sample still being classified as date driven, but appears as a necessary condition.

**Figure S7:**
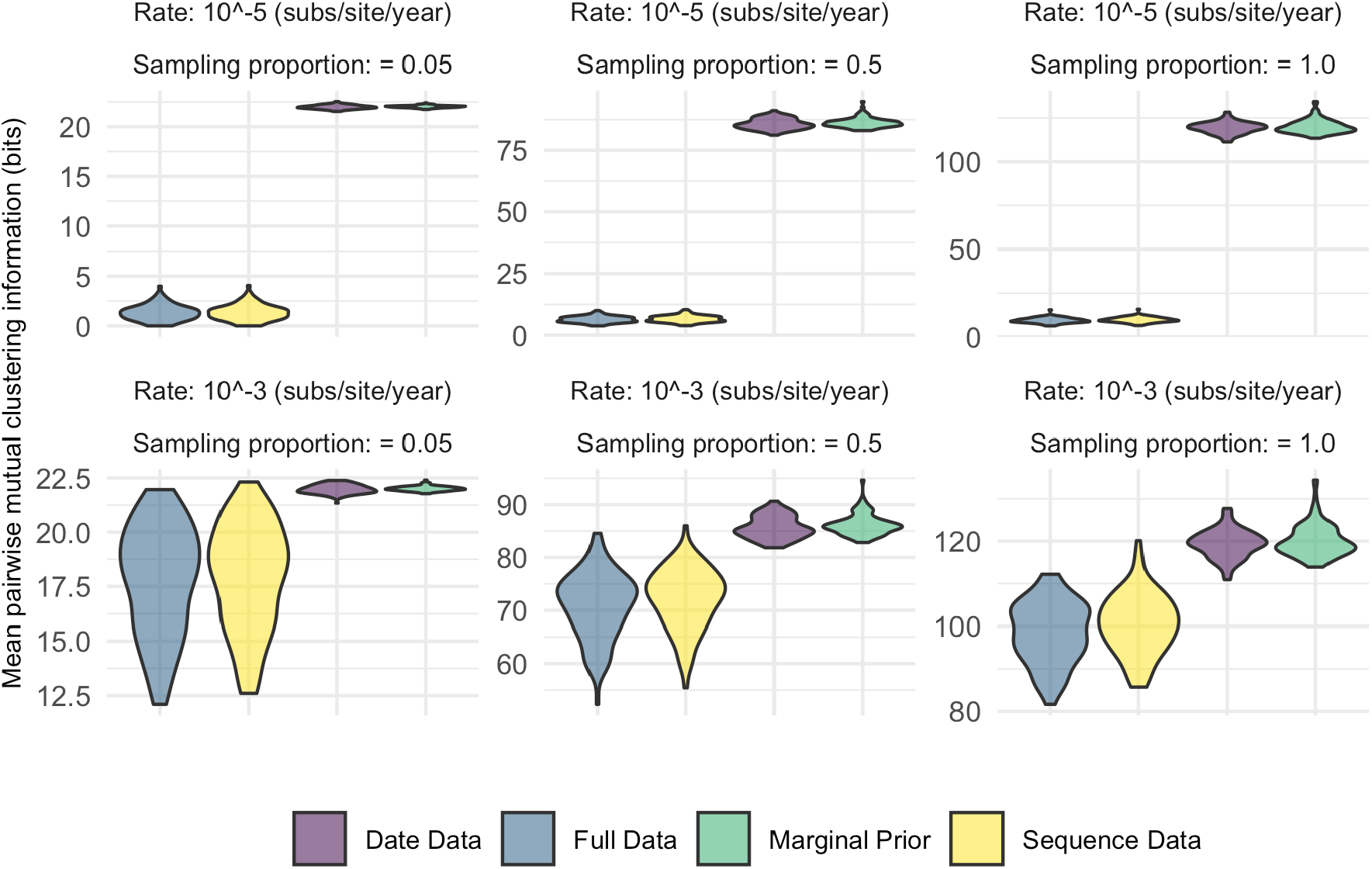
Violin plots of pairwise mutual clustering information tree difference for 100 trees sampled posterior tree distributions for each simulated dataset. 100 trees were samples from each of the 600 posterior tree distributions a total of 100 times, resulting in 60,000 datapoints here, separated by the sampling proportion and evolutionary rate used in each simulation. Uniformly, Full data has the least tree-distance between posterior trees followed by the sequence-only, dates-only, and no-data treatments.

**Figure S8:**
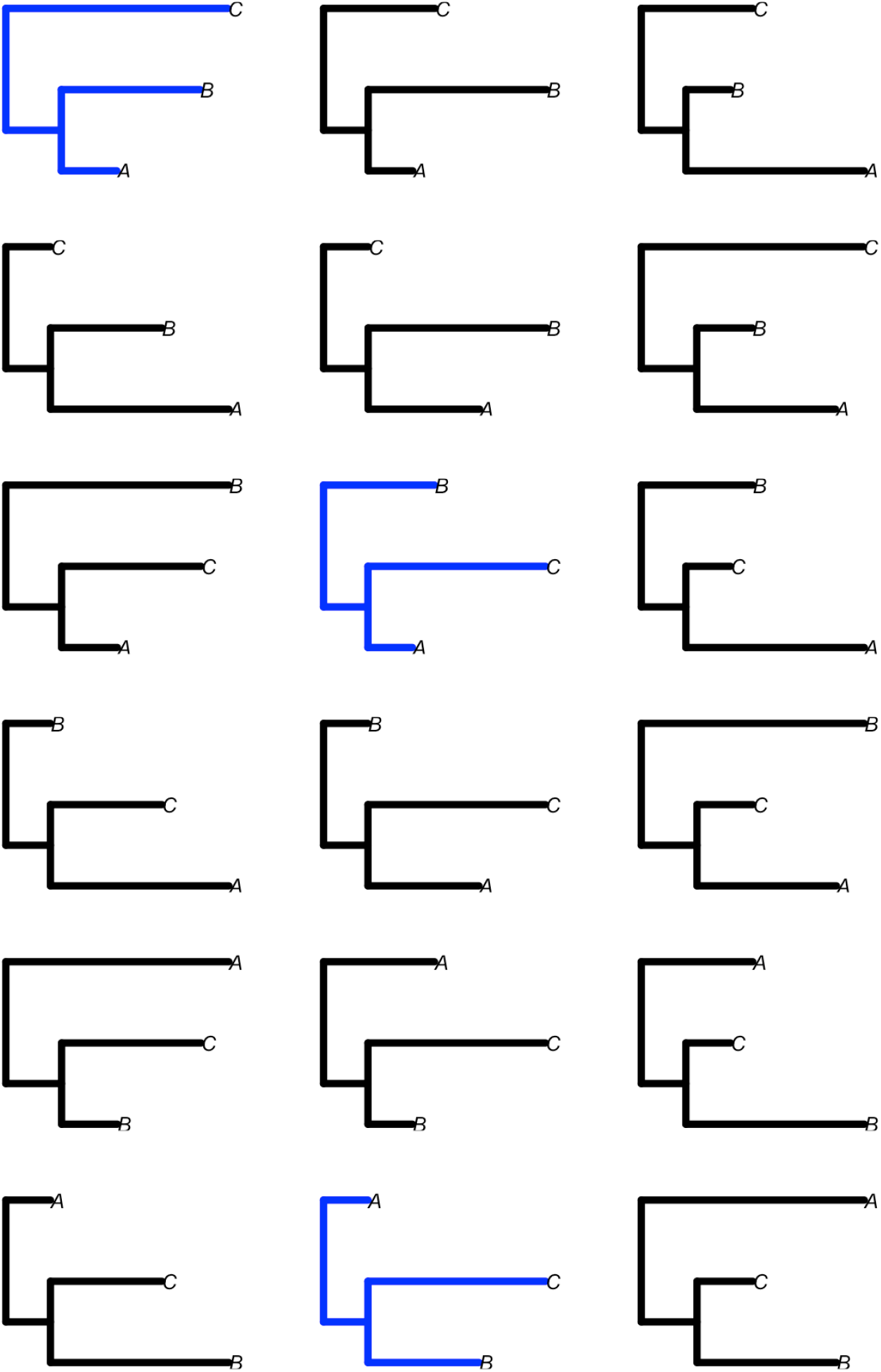
A representation of ranked tree space for a dataset of with 3 taxa. Branch lengths are in units of time. There are a total of 18 possible ranked topologies, which would be the size of tree space for an analyses of a sequence-only dataset with three samples. Trees in blue are those with the sampling chronology A, B, and then C. These represent the tree space for a hypothetical data-only dataset with three known sampling times. The key message is that tree space for sequence-only data is much larger than for date data, even with as little as three tips. Moreover, the disparity between the sequence-only tree space and date-only space grows with sample size, such that the state-space for sequence-only data is orders of magnitude larger than that for dates-only at the size of a regular phylodynamics dataset.

## Acknowledgements

LAF is grateful to Swissnex for awarding a Student Research Scholarship to foster this research. SD and LAF were received funding from the Australian Research Council (DE190100805), the Australian Medical Research Futures Fund (MRF9200006), and the National Health and Medical Research Council (APP1157586).

